# Predictive leaf metabolomics reveals systemic signatures of floral colour and form in *Camellia*

**DOI:** 10.64898/2026.01.30.702806

**Authors:** Fafa Ikram Hocini, Sylvain Prigent, Annie Lerisson, Rémy Cordazzo, Coralie Muller, Claudia Rouveyrol, Cédric Cassan, Aurélien Rey, Yves Gibon, Nicola Fuzzati, Vincent Cocandeau, Pierre Pétriacq

**Affiliations:** Univ. Bordeaux, INRAE, UMR1332 BFP, 33882 Villenave d’Ornon, France; Chanel Parfums Beauté, 93500 Pantin, France; Bordeaux Metabolome, MetaboHUB, PHENOME-EMPHASIS, 33140 Villenave d’Ornon, France

**Keywords:** Camellia, flower, machine learning, metabolome, metabolomics, predictive biology

## Abstract

The genus *Camellia* comprises more than 200 evergreen species of major economic and ornamental importance, characterised by high morphological and chemical diversity. While several species have been extensively studied for their bioactive compounds, the metabolic basis of floral trait variation across the genus remains poorly understood. In this study, a predictive metabolomics framework was applied to investigate the relationship between leaf metabolic profiles and floral traits, focusing on flower colour and floral form. Leaves from 315 individual trees, including 15 *Camellia* species and representing 1,160 samples, were analysed by untargeted metabolomics, generating a large-scale metabolic profiling dataset. A dedicated quality control strategy was implemented to ensure analytical stability across multiple injection series and flowering seasons. Penalised generalised linear models were used to uncover robust metabolic predictors associated with floral traits and to evaluate model performance through internal and external validation. Distinct sets of metabolites were associated with flower colour and floral form, with limited overlap between traits. Predictive performance was consistently higher for colour than for floral form, indicating more structured metabolic signatures for chromatic traits. The selected predictors spanned multiple major chemical classes, supporting a systemic organisation of the metabolome rather than reliance on single biosynthetic pathways. Consistently high predictive accuracies were obtained, reaching approximately 87% for both flower colour and floral form, and remaining clearly above the corresponding no-information rates (≈ 43%). Together, these results demonstrate that leaf metabolomics can be used to robustly predict floral traits in *Camellia* and highlight the potential of predictive metabolomics as a tool for early phenotype inference, quality control and selection in long-lived ornamental species.

## I. Introduction

The genus *Camellia* (Wu et al. 2022) is one of the most diverse groups within the *Theaceae* family, comprising more than 200 species of evergreen trees and shrubs mainly native to East and Southeast Asia. Widely distributed and of major economic and horticultural importance, *Camellia* species are cultivated across regions with favourable environmental conditions, although successful cultivation remains strongly dependent on mild climates and well-distributed rainfall (Jayasinghe et Kumar 2019; Zhao et al. 2021). The genus is characterised by remarkable morphological diversity, including highly variable floral traits, making *Camellia* a valuable system for studying phenotype diversity in perennial plants.

Several species are particularly prominent. *Camellia sinensis* is cultivated worldwide for tea production, one of the most consumed beverages globally (Wang et al. 2022), valued for its sensory diversity and rich content of bioactive compounds such as polyphenols and alkaloids. *Camellia oleifera* is mainly grown for the production of edible oil rich in unsaturated fatty acids (Luan et al. 2020), while *Camellia japonica* is widely appreciated for its ornamental flowers and for the high concentration of phenolic compounds found in its tissues (Pereira et al. 2022; Pereira et al. 2023). Beyond these emblematic species, however, a large part of the genus remains poorly explored, limiting our understanding of its chemical diversity and phenotypic potential.

Metabolomics is a key approach for plant studies, as it provides integrated metabolic profiles reflecting biochemical, physiological and environmental states. It enables the identification of primary and secondary metabolites involved in growth and interactions with the environment, and supports the exploration of chemical diversity and bioactive compounds for agronomic and industrial applications (Wolfender et al. 2022). In *Camellia*, the relevance of metabolomics for characterising metabolic diversity has been demonstrated through integrative analyses combining metabolomic and transcriptomic data. In *Camellia sinensis*, population-scale metabolomic profiling revealed distinct phylogenetic groups associated with characteristic metabolic signatures, including flavonoids, proanthocyanidins and alkaloids (Yu et al. 2020). These studies illustrate the capacity of metabolomics to capture structured chemical variation within the genus and to link metabolic profiles with phenotypic and evolutionary patterns.

Beyond descriptive profiling, metabolomics can be combined with chemometric and predictive modelling approaches to infer phenotypic, physiological or environmental traits. Predictive metabolomics has been successfully applied in diverse contexts, including the prediction of environmental properties such as pesticide sorption in soils (Dollinger et al. 2023), and environmental gradients or stress resilience using interspecific metabolic signatures (Dussarrat et al. 2022). At the intra-specific level, metabolomics has also been shown to predict chemical variability linked to chemotype and parental genotype, supporting its relevance for plant selection and improvement (Dussarrat et al. 2023). More broadly, integrating metabolomic data with ecological and evolutionary frameworks has revealed large-scale patterns of phytochemical diversity and hotspots of chemical richness (Defossez et al., 2021).

Despite these advances, predictive metabolomics has rarely been applied to ornamental traits, particularly in long-lived perennial species. Floral traits such as colour and form represent complex phenotypes resulting from the integration of metabolic, developmental and environmental processes. In *Camellia*, the absence of efficient functional validation systems and the long juvenile phase before flowering further limit classical experimental approaches. One hypothesis is that the leaf metabolome contains information about floral phenotypes from the juvenile stage. In this context, predictive metabolomics offers a complementary strategy to identify metabolic signatures associated with floral traits and to infer phenotypes before their visual expression. Applied to *Camellia*, predictive metabolomics offers a framework to investigate the systemic relationship between the leaf metabolome and floral phenotypes, to characterise trait-associated chemical signatures, and to develop tools for early prediction and selection. This approach directly addresses a major constraint of *Camellia* breeding and valorisation, as flowering occurs only after several years, making phenotypic screening of floral traits time-consuming and costly. By linking leaf metabolic profiles to flower colour and floral form, including multiple floral morphologies represented in this study, predictive metabolomics provides a means to anticipate floral phenotypes well before flowering. This fills an important methodological gap for perennial ornamental species, where early selection based on vegetative traits remains limited. Beyond ornamental breeding, such early predictive tools are of particular interest for applications targeting specific floral traits for industrial uses, including cosmetic and bioactive ingredient development. Overall, this work establishes predictive metabolomics as a relevant strategy to accelerate selection, reduce breeding timelines, and improve the sustainable exploitation of *Camellia* diversity.

## II. Materials and Methods

### Plant materials and sampling

Leaf samples were collected from 315 *Camellia* genotypes belonging to the Thoby botanical nursery, grown under natural conditions in Gaujacq, southwest France (43.6336° N, 0.7604° W). Sampling was performed during the flowering periods, between February-April 2023 and October-April 2024 (Table 1).

**Table 1.**
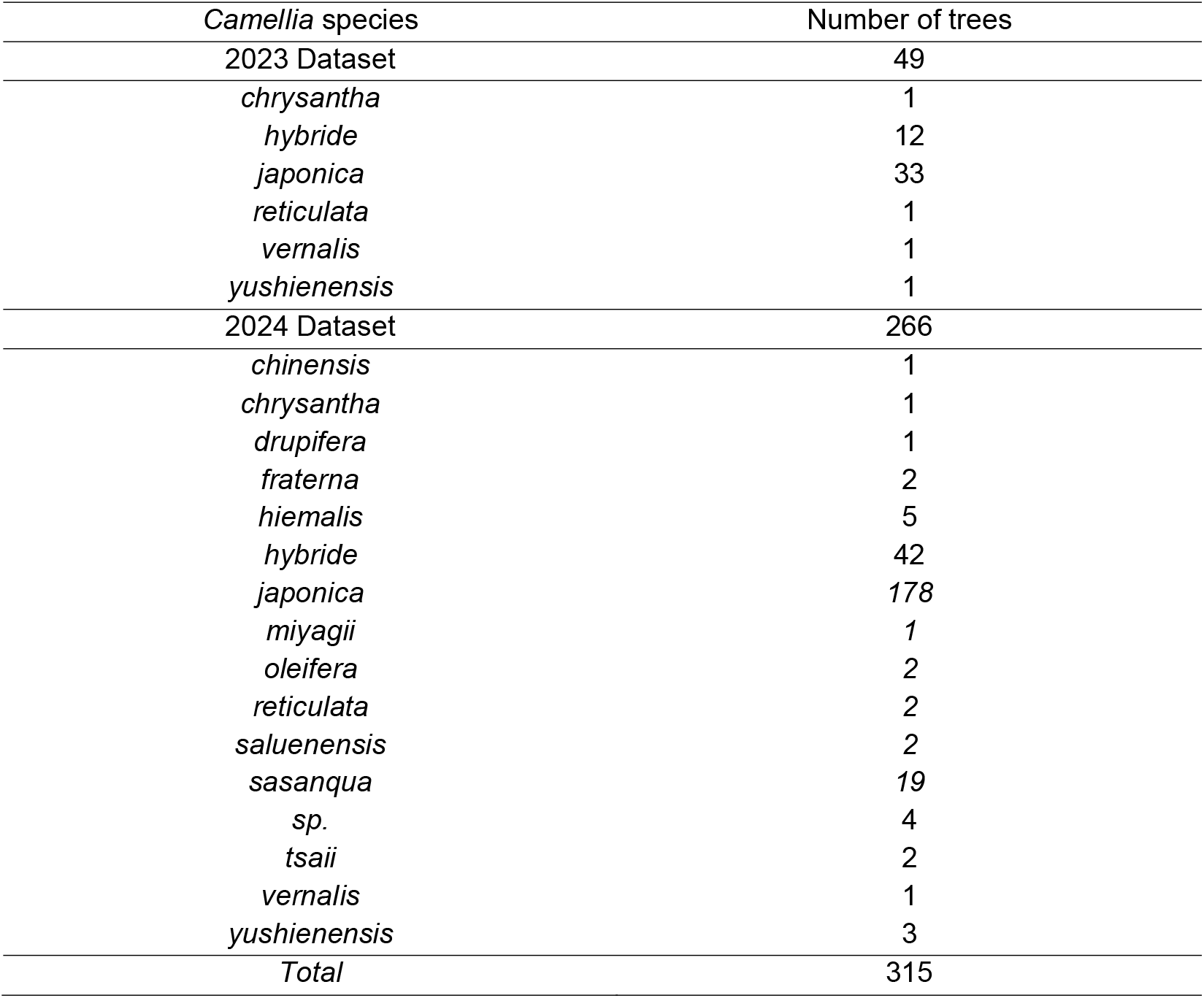
Summary of species collected in this study.

| *Camellia* species | Number of trees |
| --- | --- |
| 2023 Dataset | 49 |
| *chrysantha* | 1 |
| *hybride* | 12 |
| *japonica* | 33 |
| *reticulata* | 1 |
| *vernalis* | 1 |
| *yushienensis* | 1 |
| 2024 Dataset | 266 |
| *chinensis* | 1 |
| *chrysantha* | 1 |
| *drupifera* | 1 |
| *fraterna* | 2 |
| *hiemalis* | 5 |
| *hybride* | 42 |
| *japonica* | 178 |
| *miyagii* | 1 |
| *oleifera* | 2 |
| *reticulata* | 2 |
| *saluenensis* | 2 |
| *sasanqua* | 19 |
| *sp.* | 4 |
| *tsaii* | 2 |
| *vernalis* | 1 |
| *yushienensis* | 3 |
| *Total* | 315 |

For each genotype, fully expanded third-position leaves were collected in the morning to minimise diurnal metabolic variation. At least four biological replicates per genotype were sampled, each replicate consisting of four leaves randomly selected within the canopy. Only leaf tissues were used for metabolomic analyses.

Immediately after collection, samples were frozen on site at -28 °C, transported on dry ice, and stored at -28 °C until freeze-drying. In total, 1,160 leaf samples were obtained for metabolomic analyses.

### Sample preparation

Leaf samples were pre-ground under liquid nitrogen for 1 min 30 s at 25 Hz in a 12-position bead mill (SPEX SamplePrep, GenoGrinder 2010), with 3 stainless-steel beads of 10 mm. Powders were then lyophilised at *Bordeaux Metabolome* facility (https://doi.org/10.15454/1.5572412770331912E12) using a freeze-dryer in two programmed cycles of 42 hours (Dussarrat et al. 2022). After freeze-drying, leaf material was ground again for 1 min 30 s at 25 Hz in the same mill to obtain a fine, homogeneous powder. Flower tissues were ground in a Dongoumeau grinder for 2 min with 1 stainless-steel bead of 30 mm. Samples were processed in randomised rack positions to minimise batch effects.

### Metabolite extraction

For broad metabolite coverage, 10 mg +/-2 mg of dry material were extracted twice using an aqueous-ethanol extraction protocol (2 × 300 µL) targeting semi-polar, semi-apolar to apolar plant metabolites, as previously described (Luna et al. 2020). In brief, the extraction solvent contained 80% ethanol with 0.1% formic acid and methyl vanillate as the internal standard at 250 µg mL^−1^. Samples were extracted by sonication on ice using 300 µl of extraction solvent, then centrifuged. The supernatant was recovered, followed by a second extraction of the pellet with an additional 300 µl of solvent. The two supernatants were pooled to obtain a final volume of 600 µl of extract per sample. Extraction blanks and pooled quality controls were prepared and distributed on each rack. Tubes, plates, beads, racks, and automation equipment were as in the referenced workflow at the Bordeaux Metabolome Facility (Luna et al. 2020; Dussarrat et al. 2022).

To monitor analytical stability across extraction and injection batches, a powder-based quality control (QC) sample was prepared prior to extraction. This QC consisted of 10 mg +/-2 mg of freeze-dried leaf/flowers powder pooled from all samples, ensuring a homogeneous and representative material. The pooled powder QC was stored dry and protected from light to preserve its chemical stability over time.

For each extraction batch, five independent aliquots of this powder QC were extracted and analysed alongside the study samples. This strategy allowed batch-to-batch comparability while avoiding repeated extraction of liquid pooled extracts, which are more prone to degradation. In addition, seven extraction blanks (pseudo-extracts without plant material) were included in each batch to monitor background signals and potential contamination.

### Untargeted LC-MS-based metabolomics

Untargeted metabolomic profiling was performed using ultra-high-performance liquid chromatography coupled to a high-resolution Orbitrap mass spectrometer (UHPLC-HRMS). Analyses were carried out on an LTQ-Orbitrap Elite instrument (Thermo Fisher Scientific) equipped with an electrospray ionisation (ESI) source operated in negative ion mode.

Chromatographic separation was achieved on a Gemini C18 column (150 × 2.0 mm, 3 µm; Phenomenex) equipped with a C18 guard cartridge. The column temperature was maintained at 30 °C. The mobile phases consisted of water with 0.1% formic acid (solvent A) and acetonitrile (solvent B). The flow rate was set to 0.35 mL min^−1^. The gradient elution programme was as follows: 3% B from 0 to 0.5 min, increased to 10% B at 1 min, then to 50% B at 9 min, and to 100% B at 13 min. The column was held at 100% B until 14 min, followed by re-equilibration at 3% B until 18 min. The total run time was 18 min per injection.

Mass spectrometry data were acquired in full-scan Fourier Transform MS (FTMS) mode over an *m/z* range of 50-1500 at a resolution of 30,000 (at *m/z* 400). Data were recorded in profile mode. The ion source parameters were set as follows: spray voltage −2.5 kV, capillary temperature 300 °C, heater temperature 300 °C, sheath gas 45 arbitrary units, auxiliary gas 15, sweep gas 10, and S-Lens RF level 60%.

Data-dependent MS/MS acquisition was performed on the most intense precursor ions using higher-energy collisional dissociation (HCD). The isolation width was set to 2.0 *m/z*, with a normalised collision energy of 60%. A minimum signal threshold of 5,000 counts was applied for MS/MS triggering. Dynamic exclusion was enabled to limit repeated fragmentation of the same precursor ions. Mass calibration was performed using commercial ESI calibration solutions for negative ion mode.

Blanks and QC samples were included following a standardised injection scheme adapted to the large dataset and long analytical sequences. Instrument stability and cleanliness were ensured through weekly maintenance of the LC-MS system and chromatographic column, following the manufacturer’s recommended procedures. The column was rinsed and the ion source cleaned to limit signal drift and contamination.

For each analytical sequence, solvent blanks were injected at the beginning and at the end of the run to assess background signals and carry-over. The pooled quality control samples, prepared as described above, were injected at the start of each sequence to condition the column and then every ten samples throughout the run, allowing continuous monitoring of analytical performance and signal stability over time.

### Data processing

Raw LC-MS files were processed using MS-DIAL v4.9 (Tsugawa et al. 2019) with in-house optimised parameters. Feature detection yielded 31,410 RT-*m/z* pairs. Signal drift and batch effects were corrected using pooled biological quality-control samples, applying a LOWESS-based normalisation as implemented in MS-DIAL. In addition, methyl vanillate was added to each sample at a final concentration of 250 µg mL^−1^ and used as an internal standard to monitor consistency in extraction and injection.

Data curation removed features detected in blanks, features with a signal-to-noise ratio below 10, and features showing a coefficient of variation above 30% in pooled QC samples. After filtering, the curated dataset comprised 1,949 metabolomic features retained for downstream statistical and predictive analyses (Fig. 1). Curated metabolomics data are presented in (Supplementary Table S1), and will be made available on data.gouv.fr public repository after acceptance of this manuscript. Untargeted data were merged after sample matching, then normalised by median normalisation, cube-root transformation, and Pareto scaling using MetaboAnalystR 6.0 (Pang et al. 2024). The pre-normalised matrix and the normalised matrix are provided in Supplemental Material and will be deposited online (MetaboLight) upon acceptance of the manuscript.

**Fig. 1.**
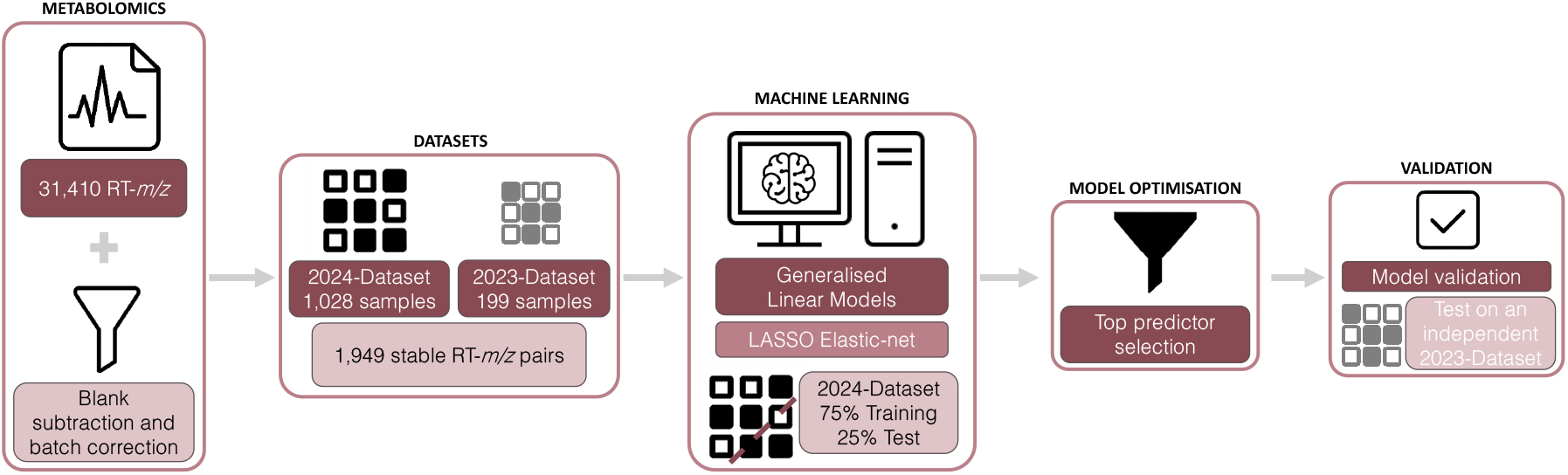
Overview of the predictive workflow. Overview of the predictive metabolomics workflow based on leaf metabolomes in *Camellia*. LC-MS profiling followed by blank subtraction and batch correction yielded 1,949 stable RT–m/z pairs. The dataset was structured into a main dataset collected during the 2024 flowering period (1,028 samples) and an independent validation dataset collected in 2023 (199 samples). Penalised generalised linear models (GLM), including LASSO and Elastic-net, were trained on the 2024 dataset using repeated random splits into training (75%) and internal test (25%) sets. Robust predictors were selected based on their recurrence across models and subsequently validated using the independent 2023 dataset.

### Statistical analyses

Principal Component Analysis (PCA) was performed in R (version 4.3.3) using the FactoMineR package (Lê, Josse and Husson 2008). The analysis was applied to the data matrix including all selected variables after normalisation, as described in the data processing section, using MetaboAnalyst v6.0. Variables were centred and scaled to unit variance, ensuring that all variables contributed equally to the analysis regardless of their original scale. Ellipses correspond to the 95% confidence regions. Hierarchical clustering and heatmap visualisation were performed in R (version 4.3.3) using the pheatmap package, with Euclidean distance and complete linkage. The resulting dendrograms were used to visualise similarities among samples and variables.

Multiclass classification analyses were performed in R (version 4.3.3) using regularised logistic regression models based on LASSO and elastic-net penalties, implemented through the glmnet (Friedman, Hastie and Tibshirani 2010) and caret packages (Kuhn 2008). Models were trained on data collected in 2024, with predictors centred and scaled to unit variance before model fitting. For model training, the 2024 dataset was repeatedly split into training and testing sets using stratified sampling on the response variable, with 75% of the samples assigned to the training set and 25% to the testing set. This procedure was repeated 100 times to account for variability induced by random data partitioning. Within each training set, model hyperparameters were optimised using 10-fold cross-validation repeated three times. A grid of 40 combinations of the elastic-net mixing parameter (α) and the regularisation strength (λ) was evaluated for each repetition. Model performance was assessed on the accuracy of the test set. To evaluate whether model predictions performed better than random prediction, classification accuracy was compared to the no-information rate, defined as the proportion of samples belonging to the most frequent class in the dataset. Predictions were considered informative when the achieved accuracy exceeded this baseline. Predictor selection was based on their frequency of inclusion in the fitted models across the 100 repetitions performed on the 2024 dataset. Variables consistently selected across multiple repetitions were considered robust predictors. To avoid information leakage, the independent validation dataset collected in 2023 was not used at any stage of predictor selection or model tuning and was employed solely for external validation. Only predictors exceeding a predefined threshold of selection frequency in the 2024 analyses were retained and used to train and evaluate models on the 2023 dataset.

### Annotation

Putative compound annotation was based on accurate mass, isotopic patterns, and MS/MS spectra computed in MSDial. Two databases were used: an in-house database of approximately 600 standard metabolites with accurate RT and *m/z* information for level and annotation, and the MetaboHUB FragHUB database (Dablanc et al. 2024), which contains a much larger set of spectral information for level 2 annotation (Sumner et al. 2007).

For metabolic network reconstruction, only robust predictors retained in at least 98% of the predictive models were considered. Annotated metabolites were mapped onto the Arabidopsis thaliana reference metabolic network using MetExplore v2.4.1.0 (Cottret et al. 2018). Mapping was restricted to metabolites explicitly present in the KEGG reference network (Kanehisa et al. 2014). Network edges correspond to curated biochemical reactions defined in KEGG and do not represent statistical correlations. The average regression coefficient across predictive models was projected onto the network nodes to visualise the relative contribution of each metabolite to trait prediction.

## III. Results

### Species and floral morphology diversity within the genus *Camellia*

To capture the floral phenotypic diversity of the genus *Camellia*, we sampled 315 individual trees (see Materials and Methods) representing over 15 species and multiple cultivars. The dataset includes all major floral forms (simple, semi-double, peony, anemone, and formal-double), and the flower colour classes: white, pink, red, and yellow (Fig. 2).

**Fig. 2.**
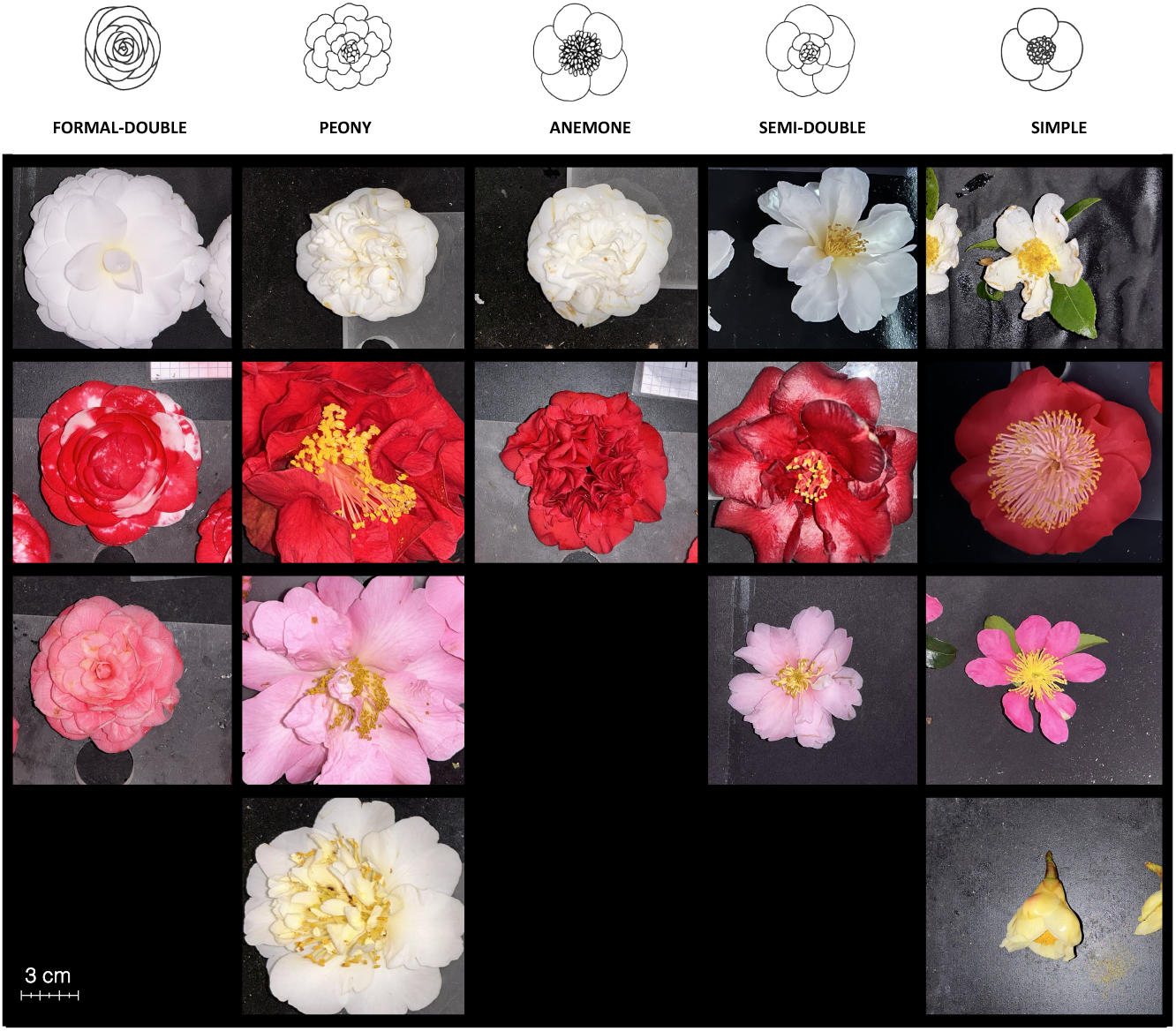
Overview of the diversity of floral forms and colours observed in this study. Representative flowers illustrating the main floral forms and colour classes included in the Camellia dataset. Floral forms comprise simple, semi-double, peony, anemone and formal-double morphologies, while colour classes include white, pink, red and yellow. Images were acquired under standardised conditions to document natural morphological and chromatic variation across species and genotypes. Scale bar: 3 cm.

Species composition and sample distribution are detailed in Table 1. The sampling prioritised broad phenotypic coverage, enabling analysis of metabolic patterns associated with flower colour and morphology across a multi-species panel. *Camellia japonica* (211 individuals) dominated the dataset, reflecting its recognised diversity in floral forms and ornamental traits (Hu et al. 2024; Fan et al. 2024; Li et al. 2025). Other species included *C. sasanqua* (19), horticultural hybrids (54), and minor representations of *C. hiemalis, C. reticulata, C. saluenensis, C. vernalis, C. yushienensis, C. chrysantha*, and *C. fraterna* (2-5 individuals each). Single individuals were sampled for *C. chinensis, C. drupifera, C. miyagii, C. oleifera*, and *C. tsaii*.

This design ensured comprehensive coverage of both genomic diversity and the full spectrum of natural flower morphologies in *Camellia*. Detailed distributions by colour, form, and species are provided in Tables S2 and S3.

Next, this unprecedented dataset of *Camellia* diversity was subjected to untargeted metabolomics, representing more than 1,600 LC-MS injections over two seasonal blooming periods (2023 and 2024). Given the large size of the dataset and the progressive addition of new samples over time, a dedicated QC strategy was implemented to support comparability across successive LC-MS injection series without systematic reinjection of all samples. For solid plant matrices, pooled QC material can be generated either by pooling extracts or by pooling sample material before extraction (Broeckling et al. 2023). Here, a pooled powder ‘biological standard’ was prepared upstream of extraction by combining identical amounts of homogenised, lyophilised material from all samples. This design provides a stable, ready-to-extract reference material that can be aliquoted and extracted per batch (Mosley et al. 2024), while enabling QC-based filtering of unstable features and monitoring of analytical performance across long sequences. After QC-based filtering for stability and reproducibility, 1,949 RT-*m/z* pairs were retained for downstream analyses.

The distribution of metabolic features across species revealed a large shared metabolomic core (Fig. 3a). For each species, the majority of detected features was common to at least two species, whereas only a small fraction was species-specific. This pattern indicated that most metabolites were conserved across the *Camellia* genus, with a limited number of features uniquely associated with individual species. A similar trend was observed for floral traits (Fig. 3b,c). Across flower colour groups, most features were shared between two or more colour classes, while only a few group-specific metabolites were detected for each colour. Red, pink and white flowers exhibited comparable numbers of shared features, whereas yellow flowers showed fewer total and specific features, consistent with their lower representation in the dataset.

**Fig. 3.**
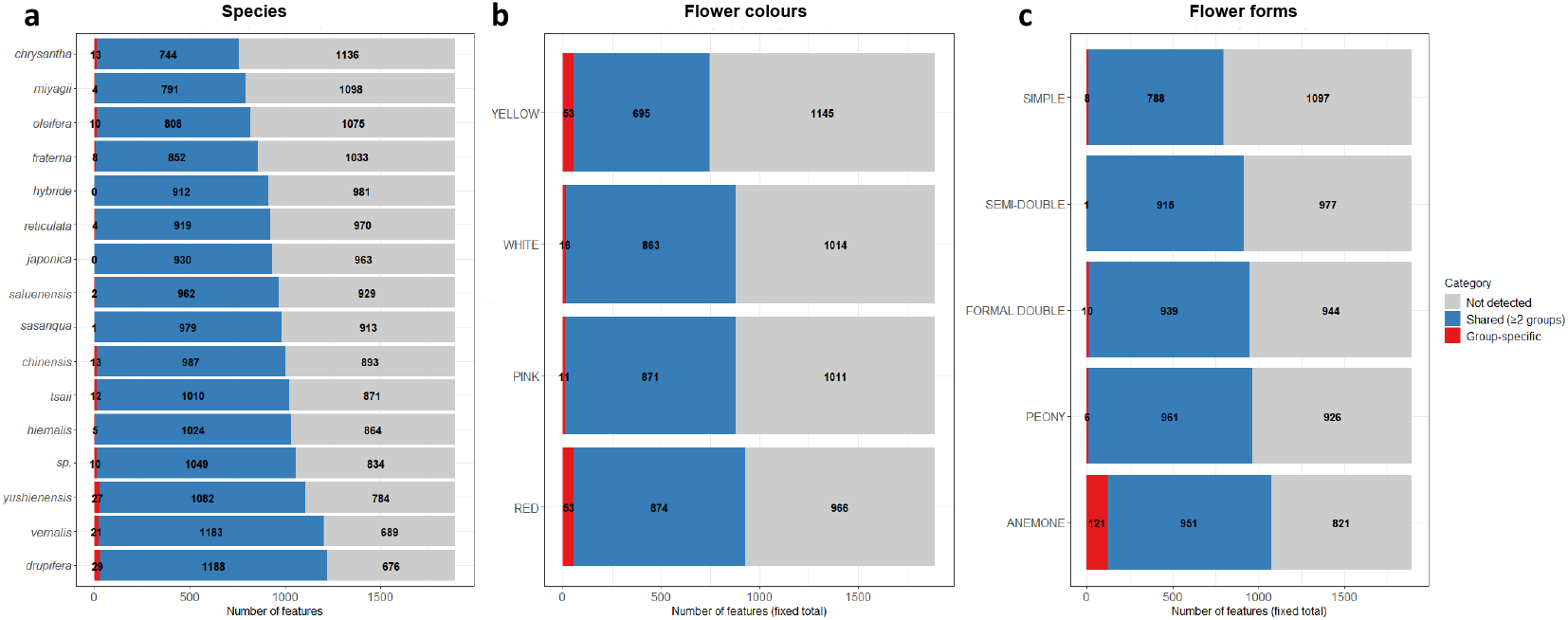
Distribution of metabolomic features across *Camellia* species, floral colours and forms. Distribution of metabolomic features detected across *Camellia* species (a), flower colours (b) and forms (c). Only features present in at least 80% of samples within each group and with an intensity above 10,000 were considered, for a fixed total of 1,949 features. Bars represent the number of features shared between two or more groups (blue), group or species-specific features (red), and features not detected in the corresponding group (grey).

For floral forms, the majority of metabolites were also shared across form classes. Only a limited number of form-specific features were detected for simple, semi-double, peony and formal-double flowers, while anemone forms displayed a slightly higher number of specific metabolites. Overall, these results indicate that both species and floral traits are embedded within a largely common metabolic background.

Principal component analysis (PCA) and hierarchical clustering analysis (HCA) revealed a partial structuring of samples according to flower colour (Fig. 4a). The first two principal components explained 16% (PC1) and 8.8% (PC2) of the total variance. Quality control samples clustered tightly in the centre of the score plots, confirming analytical stability and reproducibility across the dataset. White-flowered genotypes showed a relatively compact distribution along PC1, whereas pink- and red-flowered genotypes displayed broader dispersion, indicating higher metabolic heterogeneity. Samples corresponding to yellow-flowered genotypes occupied a more restricted region of the metabolic space. This reduced dispersion is consistent with the limited representation of yellow-flowered samples in the dataset, which mainly originate from a small number of genotypes and species compared with other colour classes (Table S2). In contrast, pink, red and white flower colours are represented across multiple *Camellia* species, contributing to increased metabolic variability along the principal components (Fig. 4b).

**Fig. 4.**
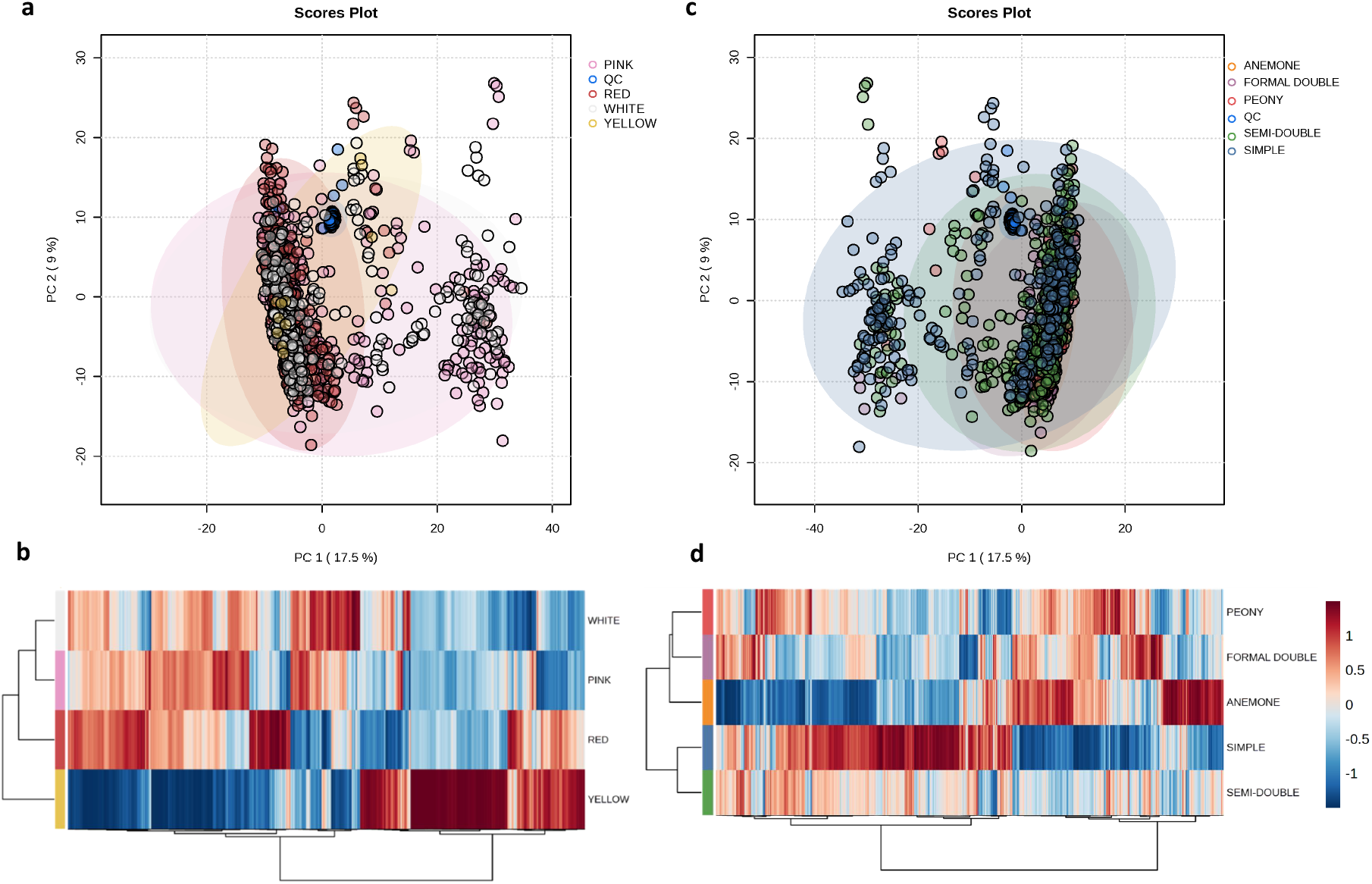
Unsupervised analysis of leaf metabolome. Principal component analysis (PCA) and hierarchical clustering analysis (HCA) of leaf metabolomic data, with samples grouped by flower colour classes (pink, red, white, and yellow; panels a, c) and floral form classes (simple, semi-double, peony, anemone, and formal-double; panels c, d). Data were preprocessed using median normalisation, cube-root transformation, and Pareto scaling. PCA ellipses represent the 95% confidence interval for sample dispersion within each colour class, while HCA displays relative abundance patterns (Pearson correlation, Ward clustering) of 1,127 and 1,473 significant metabolomic features (non-parametric ANOVA, p < 0.01, FDR-corrected) for colour and form classes, respectively. Mean metabolite intensities per class were used in HCA to improve visual clarity, and pooled quality control (QC) samples cluster tightly in PCA, confirming analytical stability and reproducibility.

A similar unsupervised approach was applied to floral form classes (Fig. 4c,d). PCA revealed substantial overlap between form classes, with no clear separation along individual principal components, reflecting the continuous and multivariate nature of floral morphology. Simple and semi-double forms exhibited higher dispersion in PCA space, whereas peony forms clustered more tightly, suggesting lower metabolic variability. Hierarchical clustering analysis (HCA) further highlighted form-dependent abundance patterns, with groups of metabolites displaying contrasting profiles across floral form classes (Fig. 4d), but without yielding discrete, well-separated clusters.

To further assess class separability, supervised PLS-DA models were constructed for flower colour (Fig. S1) and floral form (Fig. S2). While PLS-DA improved visual separation relative to PCA, substantial overlap between classes persisted in both cases, particularly for floral form. PLS-DA models showed higher performance for flower colour than for floral form. For colour discrimination, model optimisation reached R^2^ ≈ 0.62 and Q^2^ ≈ 0.53 with five components (Fig. S1d), indicating a moderate but significant predictive structure despite partial class overlap in score space. In contrast, floral form discrimination showed lower explanatory and predictive power, with R^2^ ≈ 0.44 and Q^2^ ≈ 0.30 at five components (Fig. S2d), consistent with the stronger overlap observed between form classes (Fig. S2c). Overall, these results indicate that supervised multivariate methods capture clearer metabolic structuring for flower colour than for floral form, but remain insufficient to fully resolve class boundaries, motivating the use of predictive GLM-based approaches.

Together, the predominance of shared metabolites across species and floral traits, combined with the partial separation observed in PCA, HCA and PLS-DA, demonstrates that floral phenotypes are not associated with strong isolated metabolic shifts. Instead, trait-related information appears embedded within subtle variations across a common systemic metabolome. These results indicate that unsupervised and classical supervised approaches alone are insufficient to robustly capture trait-associated metabolic signatures, motivating the use of predictive modelling strategies. Unlike projection-based multivariate methods, generalised linear models (GLM) approaches explicitly model the relationship between metabolite abundances and trait classes, allowing the identification of distributed, trait-specific predictor sets even in the presence of overlapping metabolic spaces. This strategy was therefore adopted to capture predictive metabolic signals beyond global variance structure and to assess their robustness across years and datasets.

To identify metabolites associated with floral traits, a predictive metabolomics framework was implemented (Fig. 1). Two independent datasets corresponding to samples collected during the 2024 and 2023 flowering periods, and acquired through separate injection series, were used to assess model robustness and generalisation. The 2024 dataset served as the main dataset and was randomly split into training (75%) and an internal test set (25%). This procedure was repeated 100 times, generating 100 independent models based on different random partitions of the data. Repeating the random splits allowed the intrinsic variability of the dataset to be captured and enabled assessment of the consistency with which metabolic features were selected across models.

For each iteration, LASSO and elastic-net classification models were trained to relate leaf metabolomic profiles to flower colour and floral form. Model performance was first evaluated on the corresponding internal test sets and subsequently assessed using the independent external validation dataset acquired during the 2023 flowering season. Predictor selection relied on two complementary criteria. First, the frequency of occurrence across models was used to quantify how consistently a feature was selected across repeated model runs. Second, model weights were examined to assess the magnitude and direction of each predictor’s contribution to trait prediction. Only predictors appearing in at least 98% of the models were retained as robust, resulting in 306 predictors for flower colour and 353 predictors for floral form. These reduced predictor sets were then used to build final predictive models, which were evaluated on the independent 2023 dataset (Fig. 1).

Using the 2024 dataset, high predictive performance was obtained for both flower colour and floral form, with median accuracies close to 87% for both traits (Fig. 3a). These values were significantly higher than the corresponding no-information rates (≈ 43%), which reflect the expected accuracy obtained by assigning all samples to the most frequent class given the observed class distributions. When applied to the independent 2023 dataset, model performance decreased, as expected, but remained clearly above the no-information rates. Median accuracies reached approximately 62% for flower colour prediction and 52% for floral form prediction (Fig. 5a), compared with no-information rates of about 42% and 38%, respectively. Across both datasets, colour prediction consistently outperformed form prediction, while floral form models showed a larger decrease in accuracy and greater variability. Together, these results indicate that the selected metabolic predictors capture biologically relevant information beyond class frequency alone and retain generalisation capacity across flowering seasons, with stronger robustness for flower colour than for floral form.

**Fig. 5.**
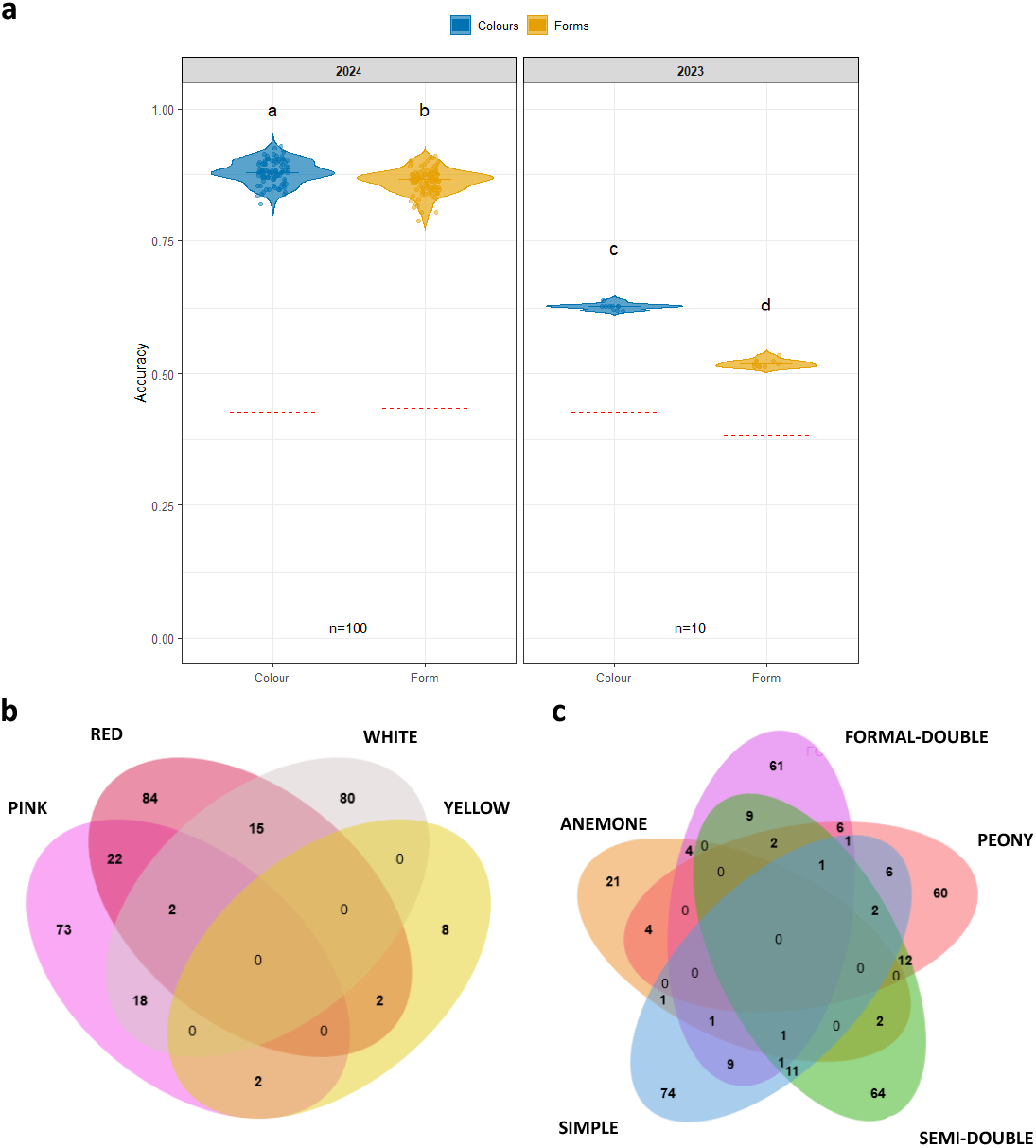
Predictive performance and trait-specific selection of metabolomic predictors for floral traits. (a) Predictive performance of linear models showing the distribution of model accuracies obtained for colour (blue) and form (yellow) predictions using the 2024 dataset (left) and the independent 2023 validation dataset (right). Violin plots summarise accuracy distributions across repeated model runs (n = 100 for 2024; n = 10 for 2023). Dashed red lines indicate the corresponding no-information rates, defined as the expected accuracy obtained by assigning all samples to the most frequent class. All pairwise comparisons were performed using the Wilcoxon rank-sum test (Mann–Whitney U test) with false discovery rate (FDR) adjustment. Different letters above violins indicate statistically significant differences between groups (FDR-adjusted p < 0.001). Bottom panels display Venn diagrams illustrating the overlap of robust predictors retained for each floral colour class (b) and floral form class (c). Numbers indicate the count of predictors specific to each class or shared between classes. Predictors correspond to features retained in at least 98% of predictive models.

### Trait-specific metabolic predictors

The composition of robust predictors retained for each trait was further examined to assess their specificity (Fig. 5b,c). For flower colour, Venn diagrams revealed that most predictors were specific to individual colour classes, with distinct sets associated with pink, red, white and yellow flowers. Overlap between colour classes was limited, and no predictor was shared across all four colours, indicating a high degree of colour specificity in the selected metabolic features.

A similar pattern was observed for floral forms. The majority of predictors were specific to individual form classes, with only minor overlap between forms. The largest sets of form-specific predictors were associated with simple, semi-double and peony forms, which are also the most represented and metabolically diverse classes in the dataset.

Overall, these results show that flower colour and floral form are associated with largely distinct and class-specific metabolic predictor sets, supporting the existence of trait-specific metabolic signatures rather than a single pool of shared predictors across floral traits.

At a global level, the aggregated representation of robust predictors (Table S4) provides an overview of the chemical space associated with floral traits in *Camellia*. This global view shows that predictive metabolites span multiple chemical superclasses for both flower colour and floral form, indicating that trait prediction relies on broad chemical diversity rather than on a single dominant class.

To refine this analysis, we focused on predictors that were unique to each trait class, excluding predictors shared between classes (Fig. S3). At this level, colour classes were characterised by distinct chemical subclass profiles. Pink-specific predictors were dominated by flavonoid-related families, including flavonoid glycosides, isoflavonoid O-glycosides, O-methylated isoflavonoids, chalcones and dihydrochalcones, together with benzoic acids and derivatives and several carbohydrate-associated features. Pink also included a small set of nitrogen-related predictors, such as purine ribonucleotides and 1-ribosyl-imidazolecarboxamides. Red-specific predictors also contained many flavonoid glycosides, but were additionally enriched in depsides and depsidones, fatty acyl glycosides, saccharolipids and cinnamic acid-related families (cinnamic acid esters, cinnamic acids), as well as linear diarylheptanoids and purine-related compounds. White-specific predictors showed a broader mix of primary- and specialised-metabolism families, including carbohydrates and conjugates, fatty acid esters, glycerophosphates, saccharolipids, triterpenoids and multiple coumarin-related families (coumarins, hydroxycoumarins, pyranocoumarins), together with several diarylheptanoid and lignan-related annotations. Yellow-specific predictors were comparatively few and mainly assigned to flavonoid glycosides, terpene glycosides, terpene lactones, sesquiterpenoids and a limited number of carbohydrate-associated features, consistent with a narrower chemical signature for this colour group.

For floral forms, trait-specific predictors also showed clear chemical subclass differences (Fig. S3). Anemone-specific predictors included flavonoid glycosides and depsides/depsidones, together with hybrid peptides, glycosylglycerols, hydroxycinnamic acid derivatives, and smaller sets of stilbenes, azaphilones and pyranocoumarins. Formal-double predictors were chemically diverse, combining flavonoid glycosides, chalcones and dihydrochalcones, linear diarylheptanoids, steroid lactones, triterpenoids, coumarin glycosides, purine derivatives and cinnamic acids, alongside several lipid-related families. Peony predictors included fatty acyl glycosides, flavonoid glycosides, chalcones/dihydrochalcones, glycosylglycerols, hydrolysable tannins, diarylheptanoids and coumarin-related annotations. Semi-double predictors were marked by flavonoid glycosides, tannins, terpene glycosides, fatty acyl glycosides, sesquiterpenoids and additional aromatic families (for instance, naphthopyranones and cinnamic acid amides). Simple-form predictors spanned flavonoid glycosides, hybrid peptides, depsides/depsidones, hydroxysteroids, glycosylglycerols, benzoic-acid derivatives and several lignan-related families. Across all classes, the model average coefficients associated with each predictor (positive or negative) indicate the direction of contribution to classification, supporting that each trait class is associated with a structured, class-specific chemical signature rather than a common predictor pool. Overall, the combination of a global chemical overview (Table S4) and trait-specific analyses of unique predictors (Fig. S3) indicates that floral colour and form are associated with structured, trait-dependent chemical signatures rather than shared or generic metabolic profiles.

### Network-based contextualisation of trait-specific metabolic predictors

To further contextualise the robust predictors, selected ions were projected onto a reference metabolic network reconstructed from KEGG pathways using MetExplore (Fig. 6). Only predictors retained in at least 98% of the predictive models were considered. Importantly, this network analysis was restricted to predictors specific to each floral trait, excluding predictors shared between colour and form as identified in the Venn diagram analysis (Fig. 5b,c). This choice aimed to highlight trait-specific metabolic organisation rather than common predictive signals. These predictors correspond to metabolites putatively annotated at level 2, based on accurate mass (MS^1^) and MS^2^ spectral information. The network was reconstructed using *Arabidopsis thaliana* as a reference biosource, and only metabolites explicitly mapped to the KEGG reference network were retained. Node colours represent the average regression coefficient across predictive models, while shaded areas indicate KEGG pathway annotations. Edges correspond to curated KEGG biochemical reactions and pathway structure rather than statistical associations.

**Fig. 6.**
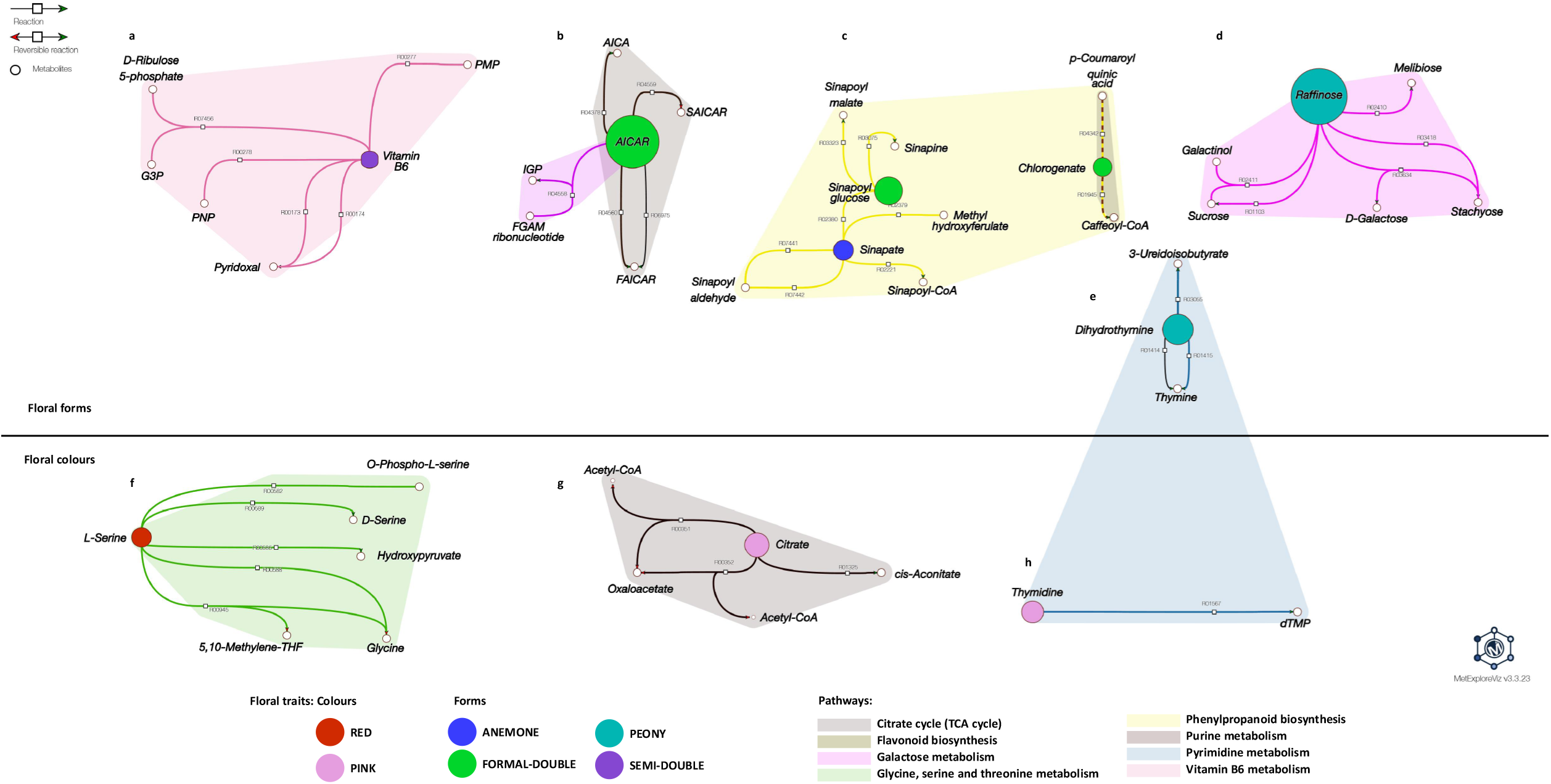
Projection of validated metabolomic predictors onto metabolic networks. Validated metabolome predictors associated with floral form (a-e) and floral colour (f–h) were mapped onto the *Arabidopsis thaliana* KEGG reference metabolic network using MetExplore v2.4.1.0. Only predictors belonging to the top 98%, as defined by their frequency of occurrence across repeated model runs, were retained for network reconstruction. Predictors were annotated at MSI level 2 based on MS^1^ and MS^2^ information and restricted to metabolites explicitly present in the reference network. Nodes represent KEGG metabolites and edges correspond to curated biochemical reactions defined in KEGG. Coloured areas indicate KEGG pathways or pathway modules. Node size is proportional to the average regression coefficient across predictive models, reflecting the relative contribution of each metabolite to trait prediction. The network visualises the distribution of high-confidence predictive metabolites across primary and secondary metabolic pathways. Reaction codes from MetExplore are displayed as white squares.

The metabolic network reconstructed for colour-associated predictors revealed a structured projection of metabolites onto the reference network. Mapped predictors were not randomly distributed but organised into a limited number of interconnected metabolic regions. A prominent region involved amino acid metabolism, centred on serine, glycine and related one-carbon intermediates, with links to folate-dependent reactions. A second connected region corresponded to central carbon metabolism, including pyruvate, acetyl-CoA and tricarboxylic acid cycle intermediates such as citrate, isocitrate and oxaloacetate. Together, these regions highlight metabolic junctions linking primary metabolism to downstream biosynthetic pathways.

In addition, a distinct region was associated with phenylpropanoid-related metabolism, including sinapoyl derivatives and related intermediates.

## IV. DISCUSSION

### Predictive leaf metabolomics captures systemic signatures associated with floral traits

This study provides evidence that leaf metabolomes contain robust and biologically meaningful predictors of floral traits in *Camellia*. The dataset assembled here represents an unusually broad survey of metabolic variability across the genus, encompassing a diversity of species and cultivated forms rarely profiled at this scale. Such breadth strengthens the generality of the predictive relationships identified and positions *Camellia* as a valuable model for exploring systemic metabolome-phenotype associations in long-lived ornamentals.

Predictive metabolomics has previously shown that complex plant phenotypes can be inferred from systemic metabolic signatures, particularly in crops such as rice (Mohanan et Kundu 2025; Riedelsheimer et al. 2012). More broadly, plant metabolomes integrate genetic and developmental information and can act as a bridge between genotype and phenotype, capturing coordinated variation across multiple pathways rather than relying on single marker compounds (Rai et al. 2025). Our results extend this concept to *Camellia*, revealing distinct sets of leaf metabolites associated with flower colour and floral form, with limited overlap between traits. Predictive performance was consistently higher for colour (Fig. 5a), suggesting that chromatic traits are linked to more coherent and stable metabolic signatures than the more developmentally complex floral architecture.

Colour-associated predictors were predominantly assigned to phenylpropanoids and polyketides, flavonoid glycosides, hydroxycinnamic acid derivatives and related secondary metabolites (Table S4 & Fig. S3). These classes correspond to well-characterised precursors and intermediates of pigment biosynthesis pathways, including flavonoid- and anthocyanin-related routes, rather than to final chromogenic compounds (Davies, Albert et Schwinn 2012; Fernie et Tohge 2017). Flower colour is indeed driven by conserved chromogenic compounds such as anthocyanins and carotenoids (Sun et al. 2025). Their prevalence in leaves is expected: pigments accumulate mainly in floral tissues, whereas upstream metabolic signatures and regulatory states are detectable systemically through untargeted LC-MS profiling. This supports the idea that the metabolic structure relevant to pigmentation is already established in vegetative tissues and later reflected in flowers.

Beyond pigment-related pathways, colour predictors also included metabolites involved in carbohydrate metabolism, lipid and lipid-like molecules, and primary metabolic intermediates. This distribution aligns with a systemic organisation of the metabolome, in which pigmentation traits reflect coordinated metabolic states rather than the activation of a single biosynthetic route (Fernie et Tohge 2017; Jiang et al. 2024). Predictors associated with yellow flowers were fewer and more chemically restricted, including terpene-related metabolites, consistent with the carotenoid-based nature of yellow pigmentation (Sun et al. 2022; Lu et al. 2025). Carotenoids are synthesised in plastids and are often weakly detected in leaf LC-MS datasets, whereas their upstream precursors and regulatory context remain measurable at the whole-plant level. However, this colour class was strongly under-represented in the dataset, and three of the five individuals assigned to the yellow group displayed very pale beige yellow shades, close to white, in contrast to two clearly yellow-flowered genotypes. This borderline phenotypic classification likely contributed to the lower robustness observed for this group, together with the limited sample size.

### Metabolic pathways, trait specificity and the resemblance between leaf and floral metabolomes

Floral traits such as colour and form are classically associated with specific biosynthetic pathways, yet they also reflect broader genome-driven organisation of plant metabolism (Davies, Albert et Schwinn 2012; Fernie et Tohge 2017). Floral organs originate from modified leaf developmental programmes, implying partial continuity in metabolic regulation between vegetative and reproductive tissues (Khojayori et al. 2024). The metabolic resemblance between leaf and flower observed here is consistent with this developmental relationship and supports the view that floral metabolic states do not arise independently but emerge from pre-existing vegetative metabolic structure.

This interpretation aligns with current understanding of metabolic evolution: plants rarely evolve entirely novel pathways, instead repurposing and modulating existing metabolic networks. The metabolic organisation captured in leaves, therefore, reflects long-established biochemical modules that underpin both vegetative and floral traits.

In this study, colour-associated predictors were not restricted to pigment end-products but corresponded mainly to phenylpropanoid derivatives, flavonoid glycosides, hydroxycinnamic acids, carbohydrate conjugates and lipid-related compounds (Table S4 & Fig. S3). These metabolites influence substrate availability, glycosylation capacity and metabolic fluxes rather than directly constituting chromogenic molecules (Sun et al. 2022; Fernie et Tohge 2017).

For floral form, predictors were distributed across a wider range of chemical classes (Table S4 & Fig. S3), and separation in multivariate space was weaker (Fig. 4c,d), reflecting the polygenic and developmental complexity of floral morphology. Floral architecture integrates multiple regulatory layers, including gene networks governing organ identity and growth (Zhao et al. 2023). Structural and regulatory genes involved in phenylpropanoid and flavonoid biosynthesis vary dynamically during flower development, indicating coordinated metabolic processes during morphogenesis (Zhang et al. 2025).

Projection of trait-specific predictors onto the Arabidopsis thaliana reference metabolic network (Fig. S3) further supports this interpretation. Predictors did not map to isolated reactions but were preferentially located at interfaces between primary and secondary metabolism, including phenylpropanoid-related modules, central carbon metabolism and amino acid-associated pathways. This organisation indicates that predictive metabolites reflect coherent metabolic regions shaped by underlying genotypic and regulatory contexts rather than direct biochemical determinants of floral architecture.

### Functional specialisation versus systemic organisation of the metabolome

Interpreting predictive metabolomics for complex traits requires distinguishing functional specialisation from systemic metabolic organisation. While a pathway-centric view assumes that phenotypes are driven by a limited number of metabolites from specific biosynthetic routes (Fernie et Tohge 2017). Plant metabolism is increasingly recognised as a highly interconnected network in which phenotypic traits emerge from coordinated variation across multiple pathways. Large-scale metabolomic studies have shown that predictive performance for complex traits often relies on multivariate metabolic signatures rather than single marker metabolites (Riedelsheimer et al. 2012; Matsuda et al. 2015).

Our results support this systemic organisation. Colour-associated predictors were distributed across several major chemical classes, consistent with the structured but partial separation observed in unsupervised analyses (Fig. 4a,b). For floral form, structuring was weaker (Fig. 4c,d), and predictors were more diffusely distributed (Table S4 & S3), reflecting the broader developmental and regulatory processes underlying morphological traits.

### Methodological limitations and perspectives

Although the predictive framework proved robust, several limitations must be acknowledged. Inferring floral traits from leaf metabolomes assumes that systemic metabolic states capture processes relevant to flower development. This assumption is increasingly supported by recent integrative metabolomic-transcriptomic studies showing that systemic metabolic regulation precedes and shapes floral differentiation (Zhu et al. 2024; Chen et al. 2025). Nevertheless, this indirect link may overlook flower-specific or transient metabolic signals, particularly for traits under strong developmental and hormonal control, as previously highlighted (Fernie et Tohge 2017; Ó’Maoiléidigh, Graciet et Wellmer 2014). Such transient signatures are often detectable only in floral tissues themselves, and their absence in leaves may limit the resolution of predictions for developmentally complex traits.

Besides, model performance was systematically evaluated against the no-information rate to account for class imbalance. Although predictive accuracies exceeded this baseline, imbalance among phenotypic classes remains a source of uncertainty, especially for underrepresented groups. This challenge is well recognised in high-dimensional omics-based prediction, where class imbalance can bias classifiers towards majority classes and reduce sensitivity to rare phenotypes (Cusworth et al. 2024; Abdelhamid & Desai 2024). Phenotypic ambiguity, such as bicoloured flowers or intermediate colour classes, may further reduce discriminability by blurring categorical boundaries, a limitation also noted in recent metabolomics-based classification studies.

Finally, analytical robustness was supported by a QC strategy based on pooled, homogenised powder prepared upstream of extraction, ensuring long-term stability across extended LC-MS acquisition series. This approach aligns with emerging consensus on best practices for QA/QC in untargeted metabolomics, which emphasises stable pooled controls, drift correction and reproducibility across acquisition batches (Mosley et al. 2024). Nevertheless, interannual environmental variation may still influence leaf metabolic profiles and contribute to residual variability, a challenge widely recognised in plant metabolomics due to the sensitivity of metabolic states to environmental fluctuations (Li-Beisson, Hirai et Nakamura 2024).

### *Camellia* as a model for predictive metabolomics in ornamental plants

Recent comparative metabolomics provides compelling evidence that floral pigmentation and other reproductive traits are grounded in metabolic architectures already established in vegetative tissues. Across angiosperms, leaves and flowers share conserved upstream phenylpropanoid and flavonoid biosynthetic modules, with precursor-enriched leaf metabolomes reflecting regulatory frameworks later redeployed in floral organs. Organ-specific pigmentation, therefore, arises primarily through differential regulation, flux partitioning and metabolite amplification rather than the invention of novel biochemical pathways (Xing et al. 2022).

This systemic continuity is consistent with the evolutionary stability of plant specialised metabolism, which is increasingly viewed as modular, flexible and evolvable. Phenotypic diversification is largely achieved through redeployment of conserved metabolic networks, particularly within the phenylpropanoid-flavonoid continuum, under the control of regulatory variation rather than pathway innovation (Wen, Alseekh et Fernie 2020). Within this framework, metabolomic profiles represent integrated functional readouts that bridge genotype and phenotype more directly than individual molecular layers.

Metabolomics-based prediction has already demonstrated utility in forecasting agronomic and developmental traits in annual crops and perennials, including yield components, quality traits and developmental outcomes (Melandri et al. 2025; Guo et al. 2022). These studies highlight the capacity of early-stage metabolic states to capture coordinated pathway activity underlying complex phenotypes.

For long-lived ornamentals such as *Camellia*, where floral traits are both economically central and developmentally delayed, the ability to infer floral phenotype from juvenile leaf metabolomes represents a substantial advance for breeding and selection. Recent metabolomic analyses in *Camellia* demonstrate strong associations between vegetative flavonoid profiles and later floral pigmentation, supporting the feasibility of systemic metabolome-phenotype prediction in this genus (Zhu et al. 2024; Zhou et al. 2025). Collectively, these findings position *Camellia* as a powerful model for exploring how conserved metabolic networks can be leveraged to predict and manipulate complex reproductive traits in perennial plants.

## V. Conclusion

Together, these results show that leaf metabolomes capture systemic metabolic states that are predictive of floral traits, reflecting underlying genotypic and regulatory organisation rather than direct causal determinants. The ability to infer reproductive phenotypes from vegetative tissues provides experimental support for the view that specialised metabolism is organised across organs and development, with floral traits emerging from pre-existing metabolic structure.

For plant biology more broadly, this work highlights metabolomics as a functional integrative layer linking genotype, development and phenotype, and demonstrates how differences in trait predictability mirror underlying metabolic and developmental complexity. By extending metabolome-based prediction to a long-lived ornamental genus, the study broadens current systems-level frameworks beyond annual crops and positions leaf metabolomes as accessible readouts of metabolic potential relevant to trait evolution and prediction across angiosperms.

## Acknowledgments

We gratefully acknowledge the Thoby nursery (Gaujacq, France) for granting access to its *Camellia* conservatory, which hosts more than 2,000 *Camellia* trees and constitutes the core biological resource of this study. We particularly thank Jean Thoby for sharing his extensive knowledge and long-standing experience with *Camellia* species, which greatly contributed to the understanding and selection of plant material. We also sincerely thank all the interns for their technical support (Alexia Travier, Haida Carmen Sane-Perez, Radia Imene Khelifi and Karim Madjid).

## Supplementary material

Supplementary Table S1. Metabolomic dataset.

Supplementary Table S2. Distribution of tree species according to colour classes.

Supplementary Table S3. Distribution of tree species according to form classes.

Supplementary Table S4. Annotation of top 98% predictors.

Supplementary Fig. S1. Multivariate PLS-DA of colour classes.

Supplementary Fig. S2. Multivariate PLS-DA of form classes.

Supplementary Fig. S3. Interactive sunburst plots of the top 98% predictor annotations.

